# A pharmacological interactome platform for discovery of pain mechanisms and targets

**DOI:** 10.1101/2020.04.14.041715

**Authors:** Andi Wangzhou, Candler Paige, Sanjay V Neerukonda, Gregory Dussor, Pradipta R Ray, Theodore J Price

## Abstract

Cells communicate with each other through ligand and receptor interactions. In the case of the peripheral nervous system, these ligand-receptor interactions shape sensory experience. In disease states, such as chronic pain, these ligand-receptor interactions can change the excitability of target neurons augmenting nociceptive input to the CNS. While the importance of these cell to neuron interactions are widely acknowledged, they have not been thoroughly characterized. We sought to address this by cataloging how peripheral cell types interact with sensory neurons in the dorsal root ganglion (DRG) using RNA sequencing datasets. Using single cell sequencing datasets from mouse we created a comprehensive interactome map for how mammalian sensory neurons interact with 42 peripheral cell types. We used this knowledge base to understand how specific cell types and sensory neurons interact in disease states. In mouse datasets, we created an interactome of colonic enteric glial cells in the naïve and inflamed state with sensory neurons that specifically innervate this tissue. In human datasets, we created interactomes of knee joint macrophages from rheumatoid arthritis patients and pancreatic cancer samples with human DRG. Collectively, these interactomes highlight ligand-receptor interactions in mouse models and human disease states that reflect the complexity of cell to neuron signaling in chronic pain states. These interactomes also highlight therapeutic targets, such as the epidermal growth factor receptor (*EGFR*), which was a common interaction point emerging from our studies.

## Introduction

Nociceptive sensory neurons are responsible for detecting changes in the environment through specific receptors and then transmitting this signal to the central nervous system (CNS) via the generation of action potentials (Basbaum et al., 2009). These nociceptors innervate almost every tissue in the body, playing a critical role in detecting injury and/or pathology to skin, joints, bones and visceral organs (Woolf and Ma, 2007; Dubin and Patapoutian, 2010). While nociceptor function is needed to navigate environments safely (Bennett and Woods, 2014) and to recover after injury (Walters, 2019), these cells can also create misery when they become persistently active (Reichling and Levine, 2009; Denk et al., 2014; Price and Inyang, 2015). Nociceptor hyperexcitability and spontaneous activity are key contributors to many chronic pain states driven by inflammation, arthritis, nerve injury, cancer or other pathologies (Woolf and Ma, 2007; Basbaum et al., 2009; Reichling and Levine, 2009; Denk et al., 2014; Price and Inyang, 2015). It is widely accepted that tissue injury is directly linked to changes in the activity of nociceptors that innervate that tissue (Price and Gold, 2018). Surprisingly, relatively little is known about the factors that are released by cells within specific tissues and how these factors act on the nociceptors innervating the tissue. Our goal was to comprehensively catalog this “interactome” because such a resource can lead to the identification of many new targets that could be manipulated to treat pain disorders.

RNA sequencing (RNA-seq) experiments have defined tissue-wide and cell-specific transcriptomes for much of the body in both mice (Usoskin et al., 2015; Schaum et al., 2018; Zeisel et al., 2018) and humans (Mele et al., 2015; Ray et al., 2018). Cell profiling experiments on normal and diseased tissues have identified key molecular players in an increasing number of disease processes (Roy et al., 2018), including disorders with a strong pain component (Hockley et al., 2018; Kuo et al., 2019). However, these studies mostly focus on gene expression within a specific tissue or across cell types in a tissue and do not characterize how multiple tissues may interact to promote disease. This type of cross-tissue interaction is especially critical to pain. Nociceptors express a wide variety of receptors that allow them to detect ligands that are produced in the tissues they innervate (Woolf and Ma, 2007; Basbaum et al., 2009; Dubin and Patapoutian, 2010). Tissue pathology frequently drives changes in gene expression resulting in *de novo* or enhanced expression of ligands (e.g. cytokines and chemokines). Since many pathological tissue states produce enhanced nociception and pain (Basbaum et al., 2009; Price and Gold, 2018; Walters, 2019), it is logical to assume that changes in ligand expression cause changes in signaling frequency or intensity through receptors expressed by nociceptors. These pharmacological interactions are prime candidates for drivers of pain states.

Here, we describe a novel computational framework that comprehensively identifies ligand-receptor mediated interactions (interactome), in a high-throughput fashion, between target tissues and sensory neurons from publicly available RNA-seq datasets. We first used this novel tool to elucidate the interactome between mouse tissues and cell-types and dorsal root ganglion (DRG) nociceptors. Then, we performed 3 case studies to demonstrate the utility of this tool for identifying potential drivers of pain states. First, we used single cell RNA-seq (scRNA-seq) data from colon-innervating nociceptors in the mouse (Hockley et al., 2018) to illustrate how these neurons interact with normal and inflamed enteric glial cells (Delvalle et al., 2018). Second, we examined how human DRG (hDRG) neurons (Ray et al., 2018; North et al., 2019) interact with macrophages taken from the joints of people with rheumatoid arthritis (RA) (Kuo et al., 2019). Finally, we assessed how pancreatic cancer (Tomczak et al., 2015) drives pharmacological interactions with hDRG neurons that could reveal new targets to provide relief for this notoriously painful disease. An intriguing theme emerging from these distinct interactomes is the prominence of epidermal growth factor receptor (EGFR) ligands as possible mediators of interactions between diseased tissue and mouse and human nociceptors. This finding is consistent with recent preclinical and clinical findings suggesting efficacy of blocking EGFR signaling for chronic pain (Kersten et al., 2013; Kersten et al., 2015; Martin et al., 2017; Kersten et al., 2019).

Genome-wide (GWAS) and transcriptome-wide (TWAS) association studies for identifying pain-related genes have been performed in multiple studies on mice (LaCroix-Fralish et al., 2011) and humans (Meloto et al., 2018; North et al., 2019) and including scRNA-seq studies (Hu et al., 2016). However, to the best of our knowledge, only one GWAS study has linked associated genes to protein signaling (Parisien et al., 2019). Our work is the first to build a scalable computational framework using RNA-seq and scRNA-seq datasets, and protein signaling pathway databases, to interrogate pharmacological interactions between cell types in health and disease with a special focus on sensory neuron related pathways. As RNA sequencing resources continue to proliferate, our tool can be used to mine for potential signaling pathways and pharmacological targets in a data-driven manner based on high-throughput assay analyses.

## Results

### Defining the cell type-enriched interactome for DRG nociceptors from 42 cell types in the Tabula Muris dataset

DRG neurons interact with nearly every tissue in the body and express an array of receptors allowing them to receive signals from distinct cell types within these tissues (Woolf and Ma, 2007; Dubin and Patapoutian, 2010). To map these potential pharmacological interactions, we curated a database of ligand and receptor pairs across the genome, based on the literature and curated bioinformatics databases (Carbon et al., 2009; Binder et al., 2014; Ramilowski et al., 2015; Yates et al., 2017; Braschi et al., 2019; Szklarczyk et al., 2019). This led to the creation of a ligand-receptor pair interactome containing more than 3000 interactions.

We first sought to examine pharmacological interactions between different classes of mouse sensory neurons and a diverse array of peripheral cell types under normal conditions. To do this, we used mouse DRG scRNA-seq data (Zeisel et al., 2018) and scRNA-seq datasets from tissues interacting with or innervated by the DRG using the Tabula Muris project (Schaum et al., 2018). While many subtypes of sensory neurons have been identified (Usoskin et al., 2015; Zeisel et al., 2018), for simplicity we clustered these into 3 well-identified neuronal subpopulations: peptidergic (PEP), non-peptidergic (NP) nociceptors and neurofilament-positive, large diameter low-threshold mechanoreceptors (NF) (Woolf and Ma, 2007; Basbaum et al., 2009; Dubin and Patapoutian, 2010). This created a broad interactome between 42 cell-types found in 19 tissues, and PEP, NP and NF sensory neurons from DRG. This establishes a ligand-receptor interaction map for sensory neurons and tissues they innervate or interact with in the mouse (supplementary file 1, sheets 1 - 42). We extracted the pairs of ligand-receptor interactions for each of these cell types where the receptor was expressed in at least one type of sensory neuron (PEP, NP, or NF) and looked for enriched pharmacology relevant gene ontology (GO) terms using Enrichr (Chen et al., 2013; Kuleshov et al., 2016). The top 5 GO terms for biological process and molecular function are shown in **Figure 1**. For the biological process GO terms found for ligand genes the “*extracellular matrix organization”* term appeared in the top 5 for all but one cell-type, and was ranked as the top 1 in 37 of the 42 cell-types. This likely occurred because among the 894 ligand genes we included in the interactome, many are secreted and 89 of them are among the 229 total genes classified under the biological process GO term “*extracellular matrix organization*”. This biases our dataset to show this particular GO term to be enriched. The *positive regulation of cell proliferation, regulation of cell proliferation, positive regulation of cell motility*, and *positive regulation of cell migration* GO terms were also enriched in both non-immune cell types and macrophages. For terms that were enriched specifically in immune cells, *cytokine-mediated signaling pathway* was enriched in most of the immune cell-types, while *regulated exocytosis* and *cellular protein metabolic process* GO terms were specific to T cells and natural killer cells (**Figure 1**). This analysis reveals that for most cell types extracellular matrix and cell adhesion ligands represent the most abundant pharmacological interaction between these peripheral cells and sensory neurons. The exception is immune cells, which primarily interact with these neurons through diffusible factors.

**Figure 1.**
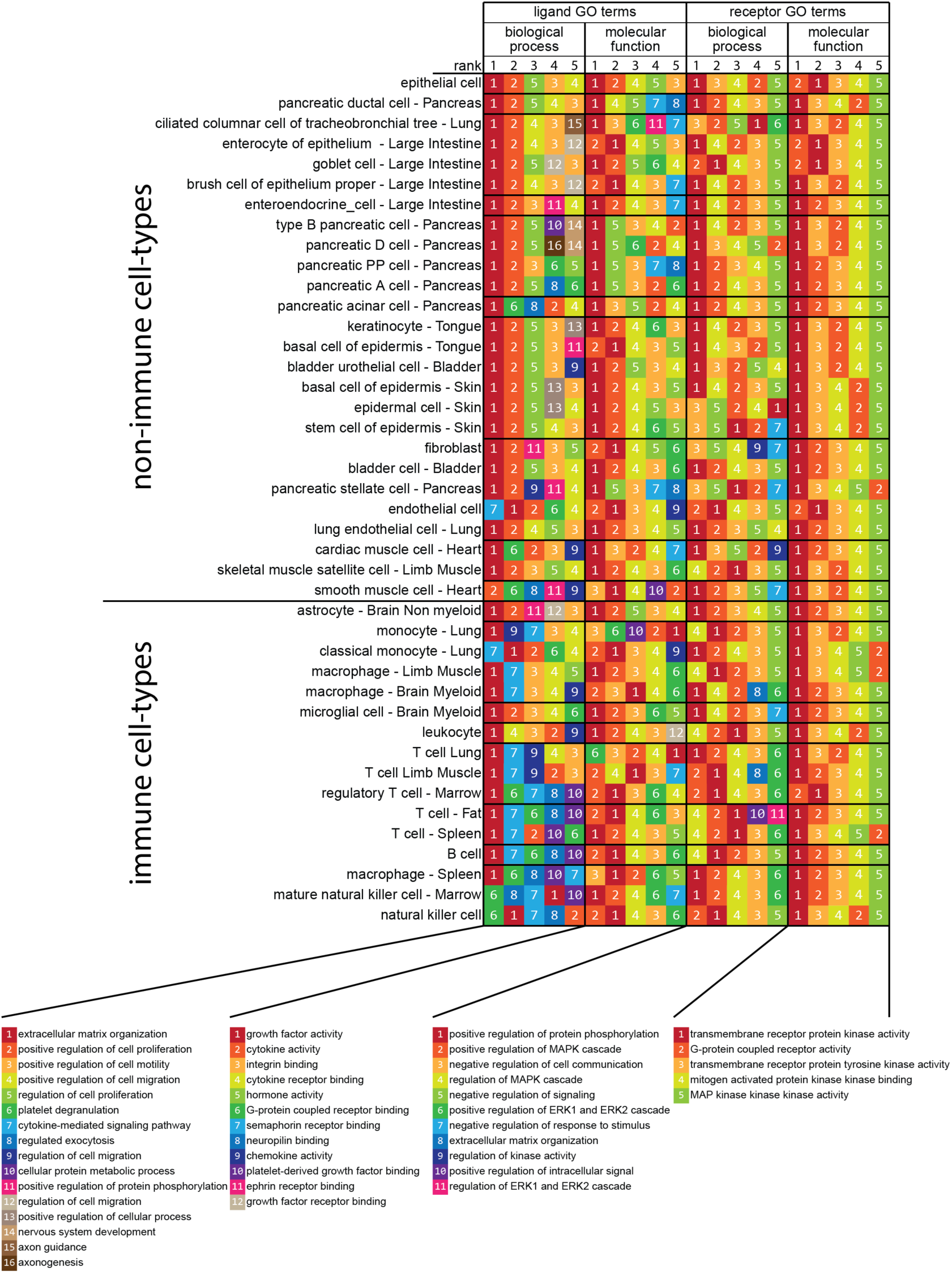
Peripheral cell type to sensory neuron interactome reveals shared principles of cell to neuron signaling across tissues. Interactome analysis was performed between all 42 peripheral cell-types from the Tabula Muris project (Schaum et al., 2018) and 3 types of sensory neurons (Usoskin et al., 2015; Zeisel et al., 2018). Only interactions where ligands were detected in the corresponding cell-type and receptors were detected in at least one of the 3 sensory neuron types were included for the GO term enrichment analysis. For the interactions identified in each cell-type, the corresponding ligand and receptor genes were separately analyzed with Enrichr for their enriched GO terms in both biological process and molecular function. The result of this analysis are shown in 4 different groups of columns. The 5 columns of color- and number-coded boxes within each of these 4 groups of columns represent the top 5 enriched GO terms in that group, ranked from left to right. Cell-types are listed as rows and ordered by the cell-type and gene bi-clustering as described in methods.

For receptor genes found in sensory neurons, the same 5 GO terms were enriched for molecular functions: *transmembrane receptor protein kinase activity, G-protein coupled receptor activity, transmembrane receptor protein tyrosine kinase activity, mitogen activated protein (MAP) kinase kinase binding, MAP kinase kinase kinase activity* (**Figure 1**). The biological process GO terms were also consistent for sensory neuron receptors identified from this interactome. The same 4 GO terms were enriched for most of the cell-types: *positive regulation of protein phosphorylation, positive regulation of MAPK cascade, negative regulation of cell communication, regulation of MAPK cascade*. Two other GO terms, *Negative regulation of signaling* and *positive regulation of ERK1 and ERK2 cascade* were enriched in non-immune cell-types and macrophages or T cells and natural killer cells, respectively. This broad interactome highlights the key role that MAPK signaling plays in transducing signals from cells throughout the body to signaling within sensory neurons. Because MAPK signaling in nociceptors plays a critical role in the generation of pain states (Dina et al., 2003; Zhuang et al., 2004; Ji et al., 2009b; Moy et al., 2017), this suggests that pharmacological interactions between nociceptors and most cell types found in the body could be capable of inducing hyperexcitability in nociceptors leading to persistent pain.

While the interactome described above shows commonalities between pharmacological interactions between sensory neurons and a variety of tissues and cells found in the mouse, it does not reveal cell type-specific interaction points that may play important roles in normal physiology and/or pathology. To find these more specific interactions in an unbiased fashion, we performed iterative hierarchical bi-clustering on cell-types and genes based on gene expression levels using Scrattch.hicat (Tasic et al., 2018). This analysis revealed 18 classes of cell types (cell clusters A-R in **Figure 2A**) and 25 gene modules spread across those cell types (**Figure 2A**). For each of these 25 gene modules we then extracted all the ligand classified genes from our ligand-receptor database for each module and constructed an interactome with receptors expressed by different classes of mouse sensory neurons (supplementary file 2). We focused on 2 module interactomes enriched in immune cell types for graphically presentation: the macrophage and leukocyte enriched cluster, and the T-cell and natural killer cell enriched cluster. We chose these based on the key role these immune cell types play in neuropathic pain models in male and/or female mice (Willemen et al., 2014; Sorge et al., 2015; Krukowski et al., 2016; Davies et al., 2019; Rosen et al., 2019; Yu et al., 2020). The ligand-receptor interactomes emerging from this gene cluster enrichment analysis revealed distinct factors expressed by these immune cells that are known to play a role in neuropathic pain states. For example, the macrophage and leukocyte cluster included *Mmp9, Il1b* and *Osm* signaling to their cognate receptors expressed by mouse nociceptors (**Figure 2B**). Macrophage recruitment by TNFα induces *Mmp9* signaling which then promotes neuropathic pain after peripheral nerve injury (Shubayev et al., 2006). *Il1b* and *Osm* have also been identified as important pain signaling molecules in previous studies in rodent pain models (Sweitzer et al., 1999) and in DRG samples from neuropathic pain patients (North et al., 2019). The T-cell and natural killer cell cluster showed many genes associated with the TNFα super-family including *Lta, Tnfsf14* and *Tnfsf11* but also highlights the *Ltbr* gene which is paired with several of these T-cell and natural killer cell expressed ligands (**Figure 2C**). This analysis also identified a specific interaction between T-cells and sensory neurons driven by interferon gamma (*Ifng*) acting through interferon gamma receptors (*Ifngr1, Ifngr2*) expressed by sensory neurons. *Ifng* had previously been found to enhance glutamate release in excitatory synapses in spinal cord and contributes to persistent pain (Grace et al., 2014). The work cited above implicates these factors in persistent pain, but our analysis shows that these immune cells express these factors at baseline. This suggests that recruitment of these immune cells to the peripheral nerve may be a key factor in driving persistent pain rather than plasticity in the transcriptomes of these cell types. This notion is supported by recent studies in rodent models (Krukowski et al., 2016; Davies et al., 2019; Yu et al., 2020) and patient transcriptional profiling (North et al., 2019). This cell-type and gene module bi-clustering approach reveals specific ligand-receptor interactions for peripheral cell types with sensory neurons that can be further mined for identification of new pain targets.

**Figure 2.**
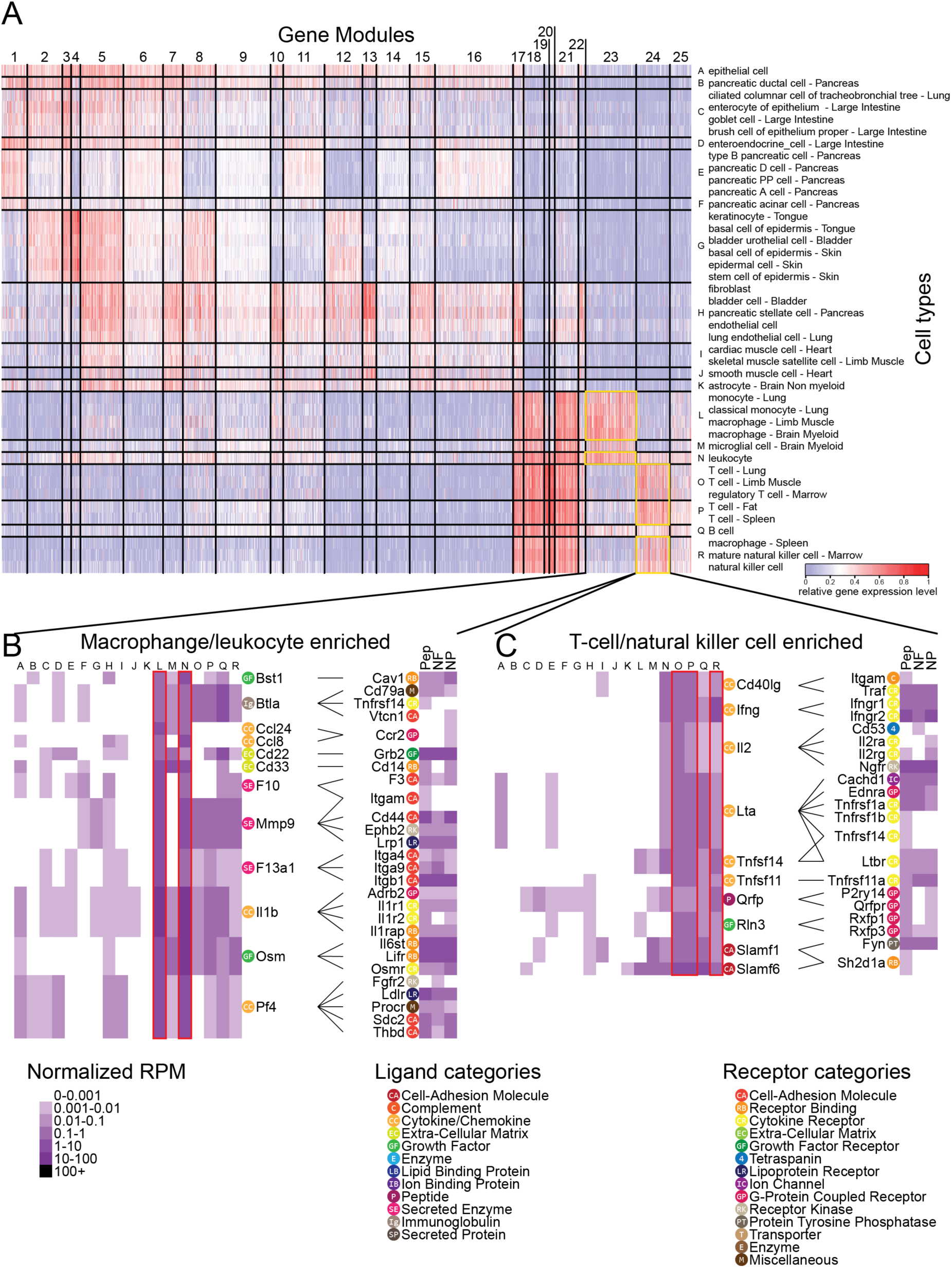
Cell type gene enrichment analysis reveals specific ligand-receptor interactions for individual cell types with sensory neurons. Iterative hierarchical bi-clustering was performed on genes and cell-types using Scrattch.hicat (Tasic et al., 2018). Eighteen cell-type modules (A, row label A-R) and 25 gene modules were identified. Two gene modules and their interactomes were chosen to be graphically presented (B, C). A) Heatmap of gene expression level with columns showing 25 gene modules identified, and rows showing 42 cell-types grouped by the 18 cell-type modules identified through bi-clustering. B, C) Heatmaps of ligand expression level across all 18 cell-type modules and receptor pair expression in 3 types of sensory neurons for each interaction are shown. Ligand genes enriched in macrophages and leukocytes are shown in (B) and ligand genes enriched in T-cells and natural killer cells are shown in (C). The cell-type modules where the ligands are enriched are highlighted with red boxes (e.g. L and N in panel (B)). The gene category shown in the legend is marked next to each ligand or receptor gene.

### Ligand-receptor interactions among cell types within the mouse DRG from the mousebrain.org dataset

The interactomes described above map ligand-receptor pairs between sensory neurons and many other cell types found in target tissues for these neurons. Sensory neurons are found within the DRG, which is composed of many different cell types, including Schwann cells and satellite glial cells. These cells are known to contribute to acute and chronic pain states (Goncalves et al., 2018; Lemes et al., 2018; Jager et al., 2020) but how they interact with sensory neurons is poorly understood. Moreover, how sensory neurons may interact with these cells through release of transmitter substances is almost completely unexplored. To examine pharmacological interactions that might occur within the DRG, we constructed ligand-receptor interactomes between NP, PEP and NF sensory neurons and satellite glial cells and Schwann cells. We did this with single cell RNA sequencing data from the mousebrain.org dataset (Zeisel et al., 2018). In our first analysis of this interactome it was clear that *Cell Adhesion Molecule* and *Extracellular Matrix* categories dominated the ligand-receptor interactions for these cell types (Supplementary file 3, sheets 1-4). This is not surprising, given the close proximity of these cells within the DRG and the obvious structural role that interactions between these cells play within the ganglion. However, because we wanted to focus on interactions driven by diffusible transmitter substances within the DRG, we chose to remove these two categories from our further analysis.

When examining which neuronal ligands potentially signal to satellite glia and/or Schwann cell expressed receptors, we made several interesting observations (Figure 3). First, growth factor interactions were the dominant category of interactions when considering neuron to glial signaling, with 30 of the 133 interactions being between growth factors and their receptors. This is consistent with previous findings in the field (Madiai et al., 2005; Yamanaka et al., 2007; Furusho et al., 2009). Second, we found indications of robust BDNF signaling within the DRG. We noted that *Bdnf*, which was expressed by NP and PEP nociceptors, has potential interactions with satellite glia and Schwann cells through its traditional receptor *Ntrk2* (TrkB), as well as through *Ddr2* (*Ntrk4*) and *Sort1*. Finally, we found that *Calca* (CGRP), a signature peptide of the PEP class of nociceptors, has an interaction with *Ramp2*, the amylin receptor in DRG glia cells. While it is known that CGRP can signal via receptors containing the amylin subunit (Hay et al., 2018), there is no previous literature on CGRP signaling through this receptor in DRG glia.

**Figure 3.**
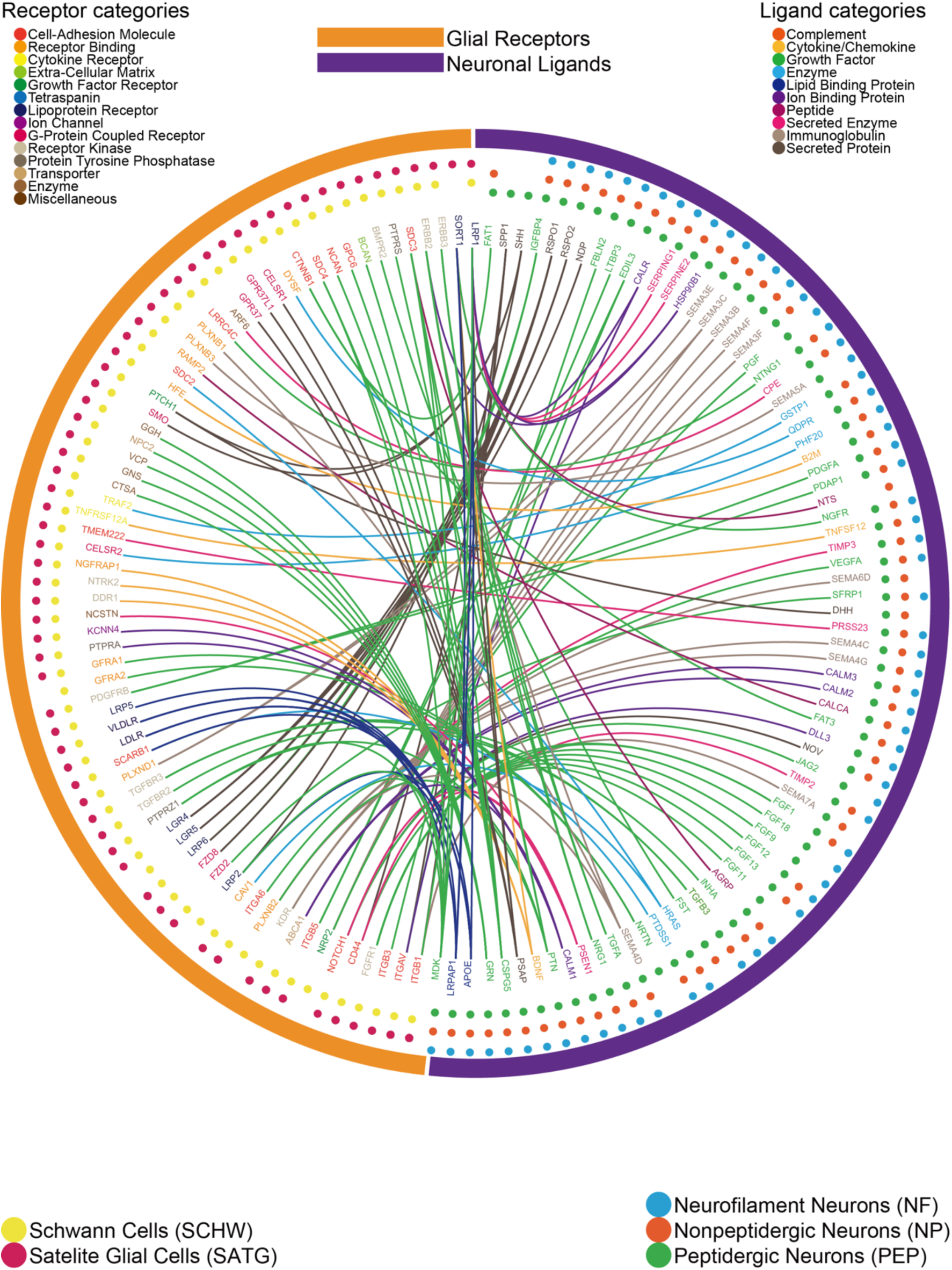
Ligand-receptor mediated interactions from neurons to glial cells within the DRG. Interactome analysis was performed to identify signaling between Satellite glia and Schwann cells and PEP, NP and NF sensory neurons. Connections with ligands expressed by sensory neurons and the paired receptor expressed by DRG glial cells are shown. Outermost circles indicate the generic class of the cells expressing the corresponding ligand or receptor genes. The middle layer shows the specific cell-type that the gene is detected in, with the dots color coded for their specific cell-types. The inner layer contains gene names, color coded for their corresponding ligand or receptor categories. Connections are marked as lines between ligand genes and receptor genes. The color of the connection line is based on the ligand gene category expressed by the neuron.

We then assessed DRG glial ligand signaling to neuronal receptors. This interactome was more diverse and revealed an increased number of these interactions. There were 133 neuronal ligands interactions with glial receptors; conversely, there were 199 glial ligand and neuronal receptor interactions. While the classes of ligands coming from glia did not fall into one main category, 56 of the 199 receptors for neurons were comprised of GPCRs, ion channels, or cytokine receptors (Figure 4). A prominent ligand-receptor interaction emerging from this dataset was the broad expression of platelet derived growth factor family genes in satellite glial cells and Schwann cells, all of which signal to a single neuronal receptor enriched in the NP class of nociceptors, *Pdgfrb*. While PDGF is known to sensitize nociceptors leading to increased mechanical sensitivity (Barkai et al., 2019; Lopez-Bellido et al., 2019), little work has been done on PDGF signaling within the DRG, making this an attractive target for further exploration. When comparing these two sets of data it is notable that there is an enormous variety in the interactions between glial ligands neuronal receptors. Our data support previous findings in the literature regarding neuronal signaling in the DRG, while identifying novel interactions which will need further investigation.

**Figure 4.**
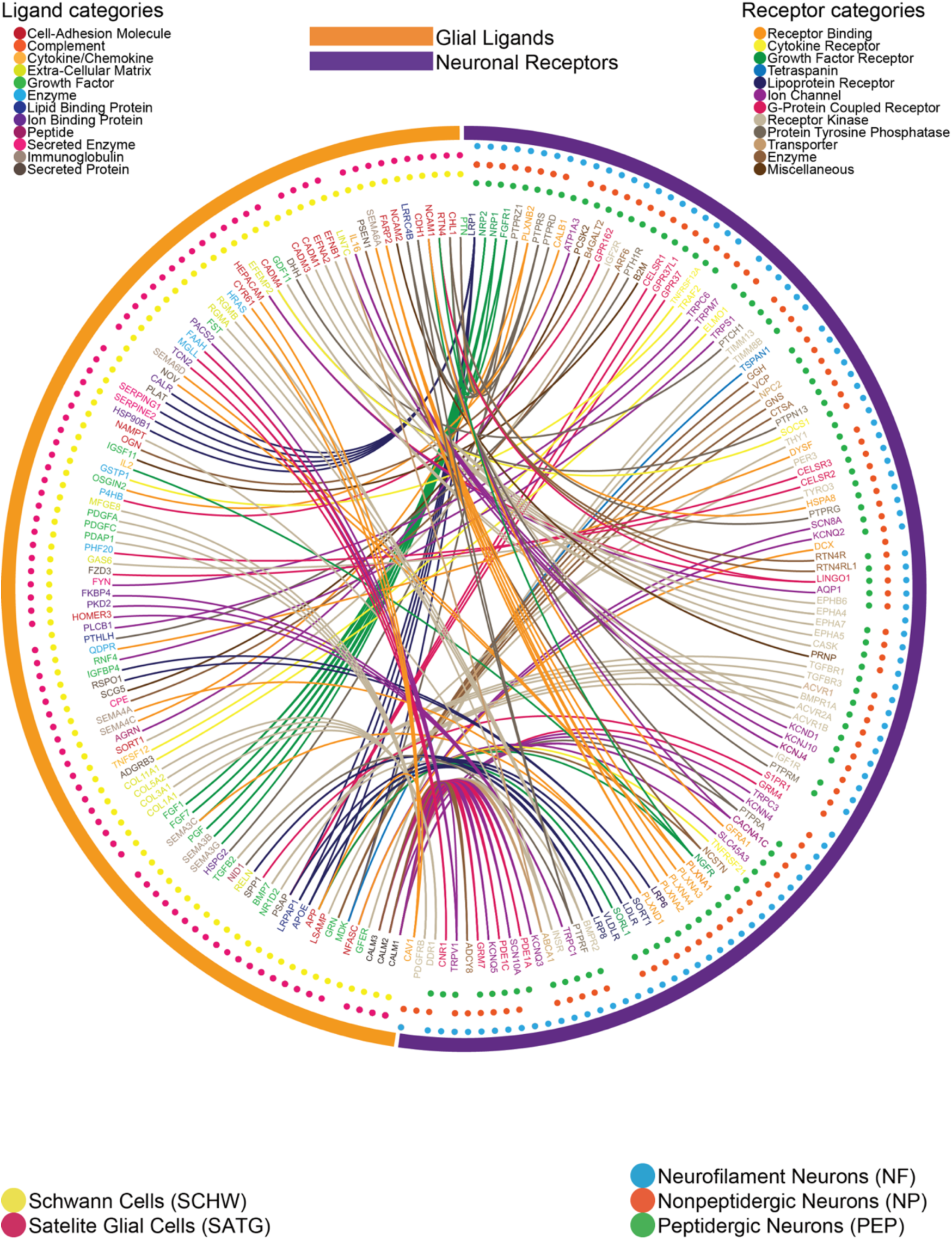
Ligand-receptor mediated interactions from glial cells to neurons within the DRG. Interactome analysis was performed to identify signaling between Satellite glia and Schwann cells and PEP, NP and NF sensory neurons. Connections with ligands expressed by glial cells and the paired receptor expressed by DRG neurons are shown. Outermost circles indicate the generic class of the cells expressing corresponding ligand or receptor genes. The middle layer shows the specific cell-type that the gene is detected in, with the dots color coded for their specific cell-types. The inner layer contains gene names, color coded for their corresponding ligand or receptor categories. Connections are marked as lines between ligand genes and receptor genes. The color of the connection line is based on the receptor gene category expressed by the neuron.

### Enteric glia to colon innervating nociceptor interactome defines gut-neuron interactions in a colitis model

Thus far, we have described interactomes between peripheral tissues and DRG neurons, as well as within the DRG, by using sequencing experiments from naïve mice. Certain pharmacological interactions may not exist in this state and may only be revealed during pathology where those interactions play a critical role in promoting disease. This principle is the basis of the use of most drugs that are used to treat disease. To explore how the pharmacological interactome of DRG sensory neurons changes in a disease state, we examined how colonic enteric glial cells react and communicate with retrogradely labelled and single cell sequenced sensory neurons innervating the colon (Hockley et al., 2018) in the mouse 2,4-di-nitrobenzene sulfonic acid (DNBS) colitis model. We used an existing dataset of RiboTag RNA-seq of enteric glial cells in this colitis model (Delvalle et al., 2018) as this technique affords cellular specificity combined with an *in vivo* inflammatory disease model. A scRNA-seq dataset of retrogradely traced sensory neurons that innervate the colon was chosen as these cells make contact with enteric glia and are at least partially transcriptomically distinct from other DRG sensory neurons (Hockley et al., 2018).

Transcriptomic changes in the DNBS treated enteric glial dataset were evaluated first. Differential gene expression analysis was performed between vehicle and DNBS treated groups to identify significantly regulated genes. These genes were then processed in our interactome analysis against retrogradely traced, mouse colonic sensory neuron scRNA-seq data. The original study identified 7 cell-types from these retrogradely traced mouse colonic sensory neurons. These cell-types were defined by expression profiles (NP, PEP, NF) and they were further defined by their anatomical location (either thoracolumbar and lumbosacral DRG, or lumbosacral only) (Hockley et al., 2018). We separated these into 5 cell-types that were found in both thoracolumbar and lumbosacral DRGs (denoted as mixed populations, m in Figure 5A), and 2 cell-types that were only found in lumbosacral DRG (denoted as pelvic populations, p in Figure 5A). The interactome between differentially expressed ligands in DNBS treated enteric glia and paired receptors enriched in one of these 7 cell-types are shown in Figure 5A (supplement table 4 sheet 1, ligand-receptor pairs that did not show any DRG neuron enrichment for any of the 7 cell types are shown in supplement table 4 sheet 2). We found that for 17 out of 22 interactions where the receptor gene was enriched in the 2 pelvic (p) specific cell-types their ligand pair gene was significantly down regulated. In contrast, in 39 out of 64 interactions where the receptor gene was enriched in mixed (m) DRG cell-types their paired ligand gene was significantly up regulated. This shows that there is a potential difference in signaling between enteric glia and colonic sensory neurons wherein inflammation relatively specifically augments pharmacological interactions between mixed population afferents whereas there is a tendency to downregulation in interactions between enteric glia and pelvic afferents (Figure 5B).

**Figure 5.**
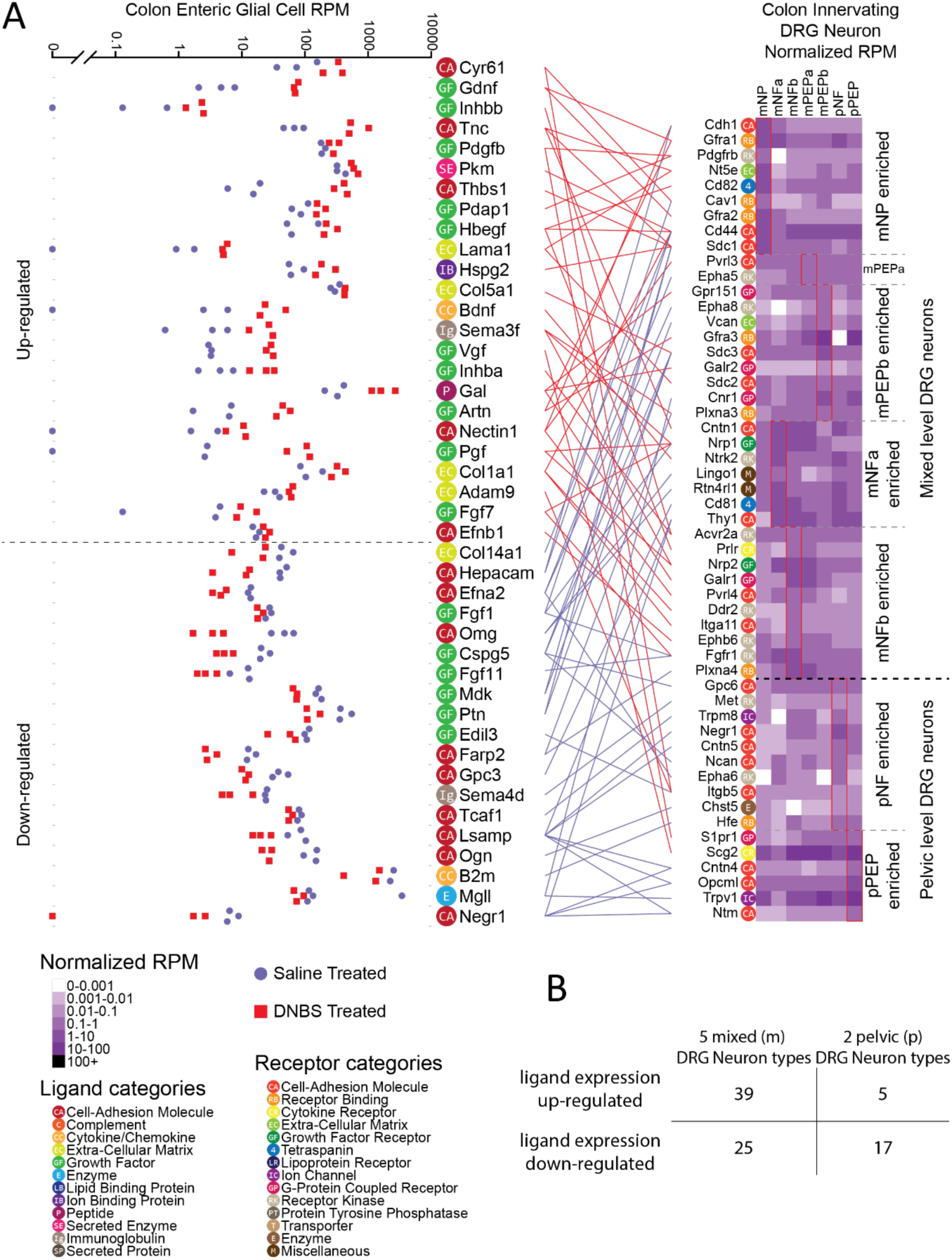
An interactome between normal and inflamed colonic enteric glia and colonic projecting sensory neurons reveals potential key drivers of visceral pain. Interactome analysis was performed to identify signaling between enteric glial cells and sensory neurons which innervate the colon. A) Connections with ligand up- or down-regulated in enteric glial cells after DNBS treatment, and the receptor expressed by DRG sensory neurons are shown. Connections are colored as red and blue, where red indicates ligand expression level was up-regulated in enteric glial cells after DNBS treatment, and blue indicates down-regulation. Ligand and receptor genes are labelled with color-coded category labels. Ligand expression level in all 6 samples (3 Saline treated (blue) and 3 after DNBS treatment (red)) are shown to the left. Receptor expression level in all 7 colon innervating cell-types are shown as a heatmap to the right. B) Table showing that up-regulated ligand genes mostly signaling to receptors found in mixed level DRG neurons.

Based on the premise that upregulated interactions may be important for pelvic pain disorders, we looked more closely at this part of the interactome. Among up-regulated ligands that signaled to DRG neurons in the mixed population we found that *Bdnf* and *Gdnf* were prominent ligands. *Bdnf* signaling to *Ntrk2* (TrkB) is known to play an important role in pain plasticity where it has primarily been studied in the context of BDNF release from primary afferents in the spinal cord (Zhao et al., 2006; Sikandar et al., 2018), but has also been linked to inflammatory visceral pain disorders (Zhu et al., 2001; Yu et al., 2012; Wang et al., 2016). *Gdnf*, which was linked to *Gfra1* and *Gfra2*, was also found to be up-regulated in DNBS treated enteric glial cells. Other studies have indicated that GDNF upregulation in target tissues enhances nociception (Albers et al., 2006; Malin et al., 2006; Queme et al., 2020). *Artn*, a ligand in the same family as *Gdnf*, was also found to be up-regulated and linked with *Gfra1* and *Gfra3*. This demonstrates a coordinated increased expression of neurotrophins in enteric glia that are likely to signal via pelvic and lumbosacral mechanisms to promote visceral pain (Zhu et al., 2001; Albers et al., 2006; Malin et al., 2006; Yu et al., 2012; DeBerry et al., 2015; Wang et al., 2016; Queme et al., 2020).

### Disease promoting macrophages from rheumatoid arthritis patients interact with hDRG through an EGR-enriched pathway

The interactome analysis described above shows that we can identify pharmacological signaling pathways in a mouse model of visceral pain. However, discoveries made in mouse models are not always consistent with actual human disease states (Bulmer and Grundy, 2011; Klinck et al., 2017). Therefore, we sought to assess whether this interactome approach could be used to identify novel targets in human disease states. This requires availability of human DRG sequencing data and sequencing data from target tissues or cells from patients with chronic pain diseases. We chose to investigate how macrophages from rheumatoid arthritis (RA) patient synovium might communicate with cell types in the human DRG, especially human sensory neurons.

A previously published scRNA-seq study of synovial tissue from RA and osteoarthritis (OA) patients identified 4 specific sub types of macrophages within the joints of patients with either of these diseases (Kuo et al., 2019). In order to find macrophage-driven interactions with human DRG neurons that are potentially responsible for promoting pain in RA, we contrasted the RA enriched macrophage cell-types with the OA enriched macrophage cell-types. Ligand genes that were highly expressed in the RA macrophages compared with the OA macrophages were selected, then filtered by whether their receptor genes were detected in human DRG RNA-seq data (Ray et al., 2018; North et al., 2019). The resulting interactome of RA-enriched macrophage ligand-receptor pairs is shown in Figure 6 (full dataset for RA and OA in supplement table 5). Interestingly, of the 20 RA-enriched ligands, 4 of them, *HBEGF, EREG, DCN*, and *HSP90AA1*, signal to *EGFR*. This suggests that the *EGFR* pathway may play a key role in promoting persistent pain in RA patients. The importance of the EGFR pathway in chronic pain has been previously noted in the literature, but not in the context of RA pain. For instance, EREG-mediated EGFR activation causes pain sensitization through PI3K/AKT/mTOR pathways in inflammatory pain models (Martin et al., 2017) and *EGFR* inhibition reduces opioid tolerance and hyperalgesia (Puig et al., 2020). Moreover, *EGFR* inhibitors have recently been used successfully for the treatment of neuropathic pain in patients (Kersten et al., 2015; Kersten et al., 2019). RA treatment has been transformed by the use of TNFα targeting biologics, but chronic pain remains a persistent problem for RA patients (Catrina et al., 2017; Krock et al., 2018). Based on our findings, we propose that *EGFR* inhibitors could potentially be repurposed to treat RA pain.

**Figure 6.**
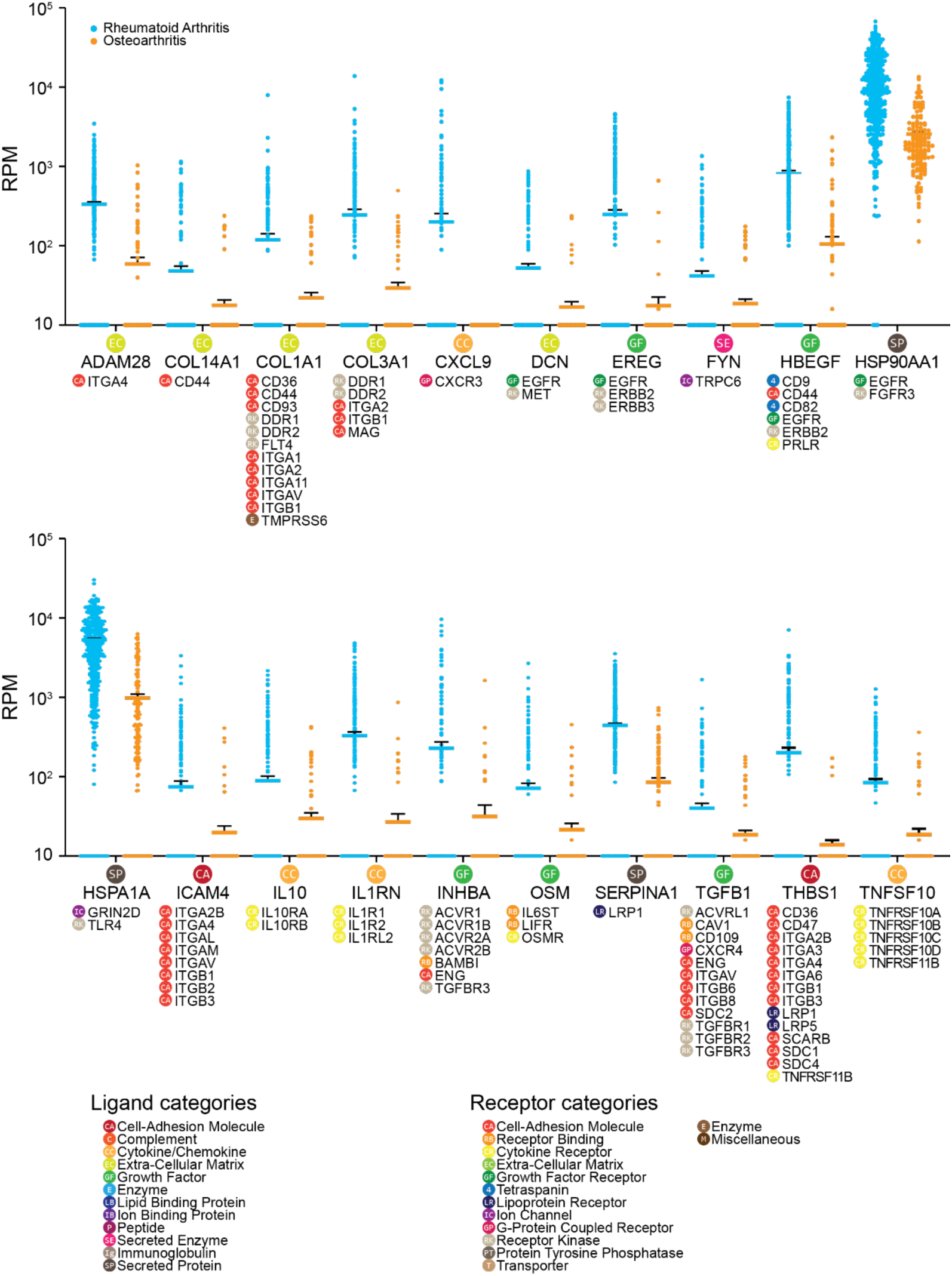
A human DRG to synovial joint macrophage interactome identifies potential drivers of persistent pain in rheumatoid arthritis. Interactome analysis was performed on differentially expressed ligand genes between synovial macrophages associated with RA and synovial macrophages associated with OA, and receptor genes from human DRG. Only ligand genes with higher expression level in RA macrophages compared with OA macrophages are shown. Gene expression level in reads per million (RPM) for each cell is shown (RA macrophages in blue; OA macrophages in orange) with the solid line as the mean and error bars as SEM. Corresponding receptor genes detected in human DRG are shown under each ligand gene. Corresponding ligand or receptor categories are color-coded and labeled for each ligand or receptor gene as shown in the legend.

### Pancreatic cancer cells suppress inhibitory and enhance excitatory signaling to human DRG neurons creating a vicious cycle for pancreatic cancer pain

We next sought to demonstrate how this interactome analysis can be used to guide identification of changes in pharmacological pathways in another human disease. Pancreatic cancer involves cancer driven mutational changes, and large scale transcriptional reprogramming. It is often associated with severe pain, and many patients are resistant to pharmacological pain treatment of any kind, requiring neurolytic treatments (Caraceni and Portenoy, 1996; Drewes et al., 2018). A better understanding of how cells from pancreatic cancerous tissue signal to DRG neurons could lead to identification of therapies that can alleviate pancreatic cancer pain.

We utilized a bulk RNA-seq dataset of pancreatic cancer tissue where we could control for individual differences in transcriptomes by having matched cancer and non-cancer pancreatic samples from each of 4 patients in the TCGA database (Tomczak et al., 2015). Ligands significantly up or downregulated in cancer samples across all 4 patients were used for the interactome analysis. These interactions were then filtered by whether their receptor genes were detected in hDRG RNA-seq data (North et al., 2019). Results of this analysis are presented in Figure 7 and 8. Among 41 ligand-receptor pairs identified, we noted that certain mediators that are well-known pain suppressing ligands showed decreased expression in cancerous tissue compared with healthy tissue. These included the endogenous opioid ligand *POMC* and the anti-inflammatory cytokine *IL10*. On the other hand, expression of many pain promoting and/or inflammatory ligands was increased, including *SHH, TGFA, TFF1*. These findings suggest that a central problem in pancreatic cancer pain may be a loss of balance between pain suppressing the pain promoting signaling that is found within the normal pancreas. Notably, 4 of these 41 ligands, *CEACAM1, FGF1, TFF1, TGFA*, are known to signal through *EGFR*, suggesting that the *EGFR* pathway may play an important role in driving pain in pancreatic cancer patients. While these *EGFR* ligands have been previously studied in the context of cancer, where *EGFR* is a well-known target, they have not been widely studied in the context of pain. Interestingly, *EGFR* inhibitors have been shown to provide pain relief in previous cancer clinical trials (Bezjak et al., 2006).

**Figure 7.**
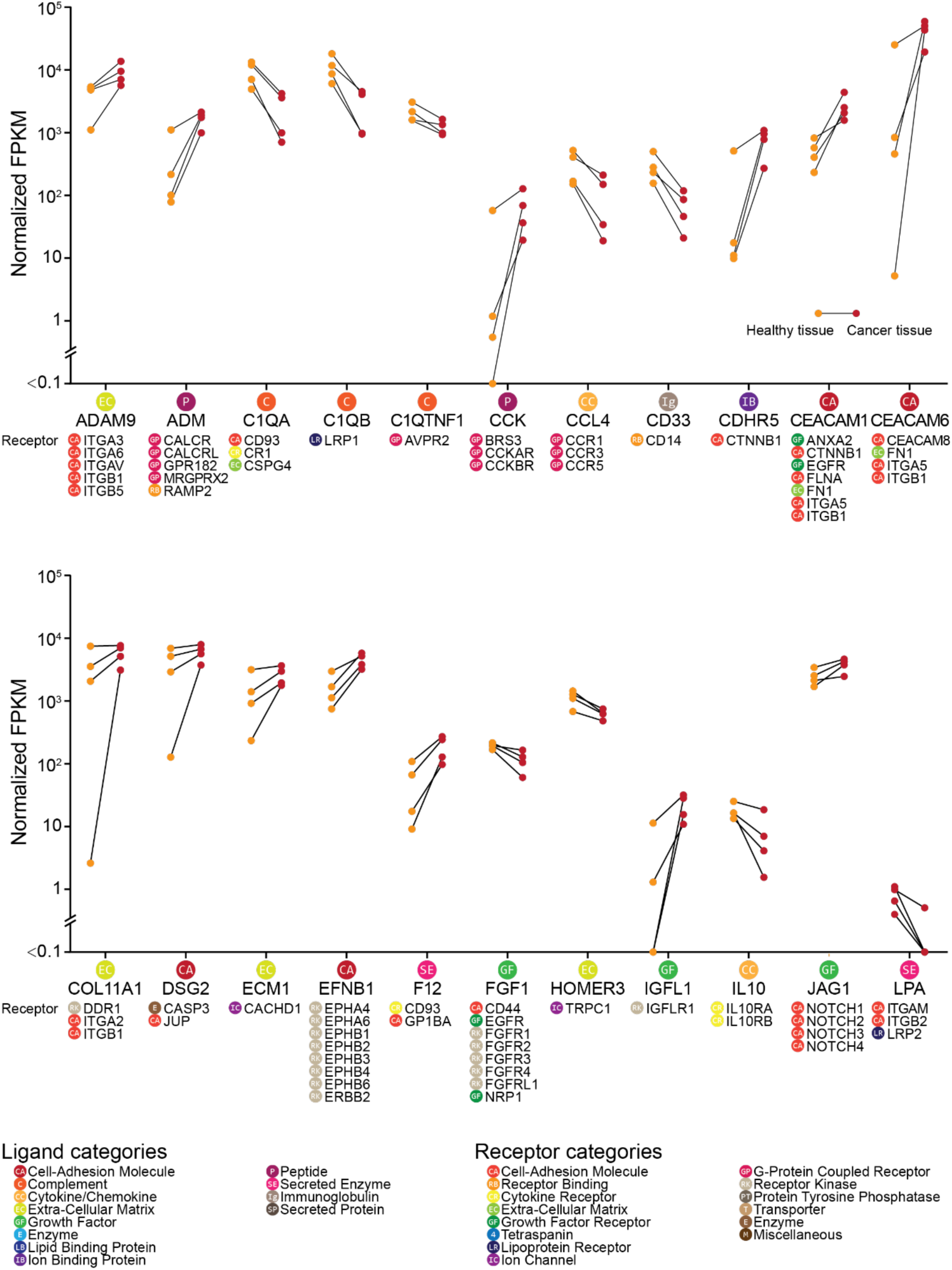
An interactome between normal and pancreatic cancer tissue in humans identifies potential pathways for treatment of pancreatic cancer pain (ligands A – L). Interactome analysis was performed on differentially expressed ligand genes between healthy and cancerous tissue from 4 individuals with pancreatic carcinoma. Only ligand genes with a corresponding receptor gene expressed in the human DRG dataset are shown. Gene expression level in fragments per kilobase million (FPKM) is shown (healthy tissue in orange; cancerous tissue in red) with the connected line marking the samples from the same patient. Corresponding receptor genes detected in human DRG are shown under each ligand gene. Corresponding ligand or receptor categories are color-coded and labeled for each ligand or receptor gene as shown in the legend.

**Figure 8.**
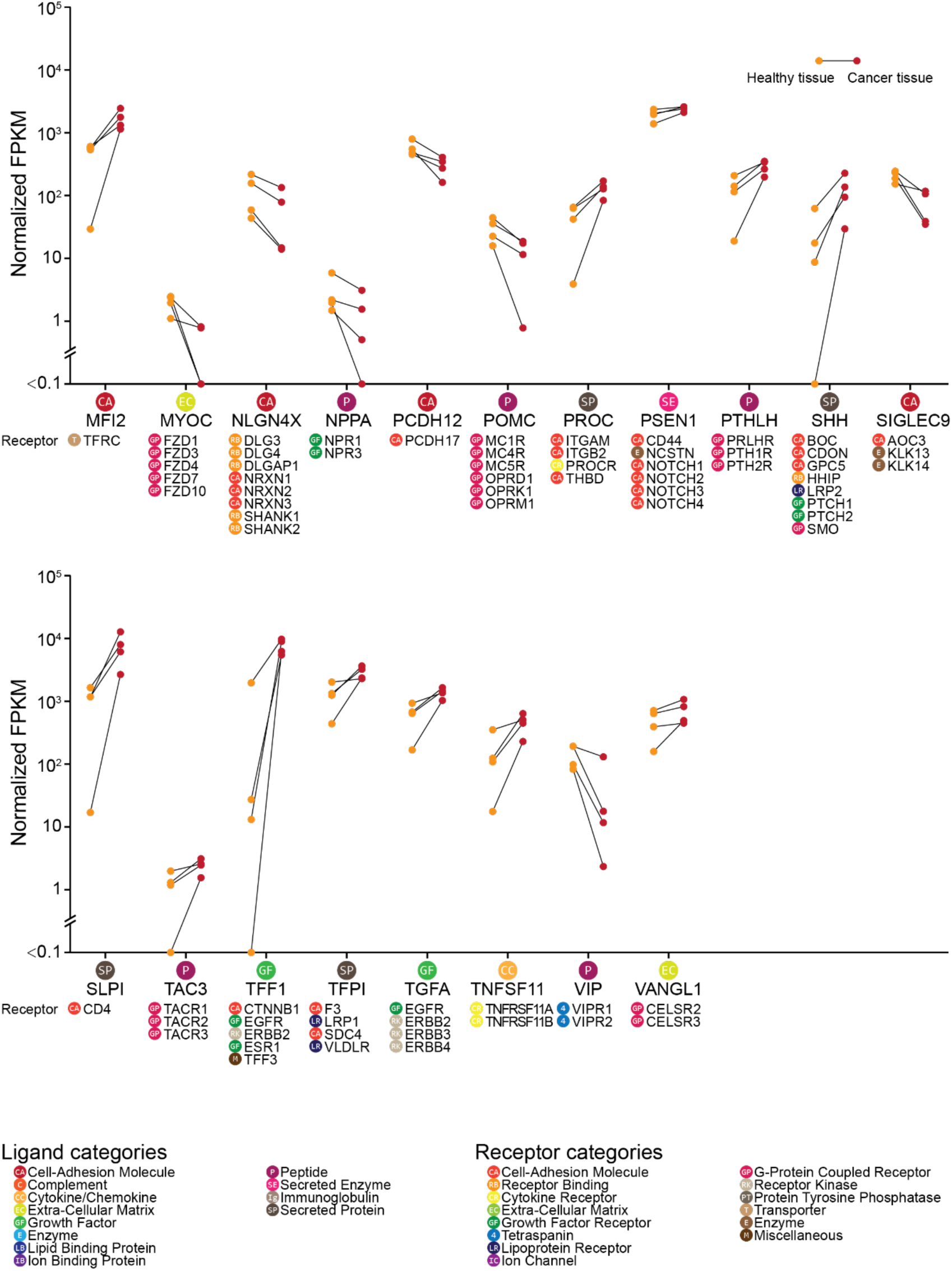
An interactome between normal and pancreatic cancer tissue in humans identifies potential pathways for treatment of pancreatic cancer pain (ligands M – V). Interactome analysis was performed on differentially expressed ligand genes between healthy and cancerous tissue from 4 individuals with pancreatic carcinoma. Only ligand genes with a corresponding receptor gene expressed in the human DRG dataset are shown. Gene expression level in fragments per kilobase million (FPKM) is shown (healthy tissue in orange; cancerous tissue in red) with the connected line marking the samples from the same patient. Corresponding receptor genes detected in human DRG are shown under each ligand gene. Corresponding ligand or receptor categories are color-coded and labeled for each ligand or receptor gene as shown in the legend.

## Discussion

We have created an interactome framework for examination of how specific subtypes of cells in the body interact with sensory neurons that innervate the target tissues where these cells reside. This resource can immediately be used to mine interactions between sensory neurons and many of the cell types found in the bodies of mice. Many of these ligand-receptor interactions are generic, however, a surprisingly large number of them show remarkable specificity. For instance, T-cells and natural killer cells appear to use lymphotoxin alpha (*Lta*) to lymphotoxin beta receptor (*Ltbr*) as a unique mechanism to signal to nociceptors. Our work also elucidates how ligand-receptor interactions change in chronic pain disease states such as rheumatoid arthritis and pancreatic cancer. This computational method can be useful for identifying new targets for disease treatment (e.g. EGFR inhibitors for rheumatoid arthritis). We anticipate that continuing advances in sequencing techniques (Stark et al., 2019), such as spatial transcriptomics (Lein et al., 2017), and their application to human disease tissues will enable targeted therapeutic discoveries using this interactome framework.

One of the key findings emerging from our work is the complexity of the potential ligand-receptor interactions that are found in these interactomes. Pain is widely acknowledged to be a complex disease, but most pain therapeutic development focuses on a single factor, such as NGF or CGRP sequestering antibodies, or receptor or enzyme antagonists (Woodcock et al., 2007). Some of these approaches, for instance NGF (Brown et al., 2012) and CGRP (Sun et al., 2016) targeting, have been effective in the clinic. However, not all patients respond to these therapeutics and even when patients do respond, these therapies are not cures. Our work shows that chronic pain disease states are accompanied by complex changes in ligands produced in diseased tissues and that many of these ligands have considerable potential to have an action on the nociceptors that innervate that tissue. This likely means that multiple ligand-receptor interactions need to be simultaneously targeted to effectively treat chronic pain states. Of course, this is not a new concept, but our work starts to provide a toolkit to quantify these ligand-receptor interactions and design therapeutic strategies that have an increased chance of success. Continuing to develop transcriptomic maps of human tissues, at the bulk and single cell level, including in disease states, will ultimately be needed to achieve this goal. Such efforts are well underway and the technology to do such studies at the individual patient level are rapidly becoming available (Stark et al., 2019).

Another important finding is the degree to which ECM and cell adhesion molecules govern ligand-receptor interactions between peripheral cells and sensory neurons. These interactions dominated our ligand-receptor interactomes and many of these interactions have not been studied at all in the context of sensory neurobiology. Several studies in the past decade have pointed out the key role that ECM molecules play in the development of chronic pain states (Ji et al., 2009a; Reichling et al., 2013; Parisien et al., 2019), but, again, these studies have only focused on a small number of the many interactions that were apparent in our interactomes. These ECM and adhesion molecule interactions may also play a critical role in recruitment and proliferation of immune cells to peripheral nerves and the DRG after nerve injury. Insofar as these neuro-immune interactions in the periphery are emerging as key players in nerve regeneration (Davies et al., 2019) and neuropathic pain (Ji et al., 2016; Yu et al., 2020), gaining a better understanding of how this occurs will yield new insight into disease states. Therefore, this is almost certainly an area that is ripe for further exploration from the perspective of fundamental neurobiology knowledge and therapeutic target discovery. Our datasets lay a foundation for further exploration along these lines.

A theme emerging from our interactomes built using sequencing data from human disease was the involvement of EGFR ligands and the EGF receptor. Previous studies have implicated the EGFR pathway with chronic pain. Genetic associations studies link the EGFR and the EGFR ligand EREG chronic temporomandibular joint pain (Martin et al., 2017). Animal pain models suggest that EGFR activation by epiregulin encoded by the EREG gene promotes inflammatory and neuropathic pain and that EGFR signaling is critical for pain promoting effects of opioids (Martin et al., 2017; Puig et al., 2020). Finally, several clinical trials have been done with EGFR inhibitors for neuropathic pain and some of these have been positive (Kersten et al., 2013; Kersten et al., 2015; Kersten et al., 2019). Our results point to a diversity of EGFR ligands that are upregulated in painful tissues such as joints of people with rheumatoid arthritis and in pancreatic cancer. These ligands were distinct in these clinical cohorts, and did not include EREG, which is reported to be unique among EGFR ligands in sensitizing nociceptors in mice (Martin et al., 2017). Given that these different ligands may have differential signaling bias when activating EGFR, it is likely necessary to study their effects on human nociceptors as several published reports have demonstrated important differences between rodent and human nociceptors (Davidson et al., 2016; Rostock et al., 2018; North et al., 2019; Moy et al., 2020). Nevertheless, our findings are promising from the perspective of broadening study of EGFR inhibitors for different chronic pain conditions.

There are some important limitations to our work. The first is that most of the interactomes presented are not precisely matched for the innervation of the target tissue. The exception is the colonic sensory neuron to enteric glia interactome. Future work will focus on building additional interactomes where the target cell types are matched to the subsets of nociceptors that specifically innervate those cells and, if possible, taken from samples with the same pathophysiological state. This will have important implications for more precise target identification. The second is that single cell resolution is not yet available for the hDRG so the human disease-based interactomes should be interpreted with some caution. Our bulk sequencing data affords a broad view of possible interactions between target tissues and hDRG neurons, but single cell sequencing on hDRG neurons would improve confidence in targets emerging from these types of experiments. A final limitation is in gaps in the ligand-receptor interactome. While we have made a concerted effort to include as many enzyme-derived small molecule interactions as we can in this database, it is not comprehensive. There are also many ligand-receptor interactions that are not yet known, and these are necessarily not part of our interactome. Therefore, while our work elucidates many aspects of ligand-receptor interactions that are potentially involved in driving painful disease states, it cannot be viewed as a comprehensive resource.

The work presented here provides an unprecedented view into how different somatic cells interact at the pharmacological level with the sensory neurons that innervate the entire body. Our findings create an important resource for the field, and elucidate new therapeutic targets for several chronic pain states. Our interactome framework is scalable for rapid advances in RNA sequencing datasets and technologies (including other high throughput transcriptomic or proteomic assays) that are happening in nearly every aspect of biological sciences and medicine. Based on the goal of the research, our generated interactomes can be ranked by a variety of criteria like disease association scores, tissue specificity of expression, and specificity of ligand-receptor interaction making our tool relevant in a wide variety of clinical and biomedical research settings. We envision this tool to be useful in identification of new therapeutic targeting strategies for specific chronic pain disorders and an important step toward personalized medicine strategies that can be applied at the transcriptomic level.

## Materials and Methods

### Mouse Sequencing Resources

#### scRNA-seq from 42 cell-types from Tabula Muris

The Tabula Muris dataset (Schaum et al., 2018) contains scRNA-seq data generated by 2 different methods: Smart-seq2 sequencing of FACS sorted cells (FACS method) and microfluidic emulsion method. The scRNA-seq data generated by the FACS method includes more tissue-types and has a higher number of genes detected per cell, therefore we used that data for our analysis. From all cell types identified in the Tabula Muris dataset we selected cells from tissues that are strongly innervated by DRG neurons and from those we chose the 42 cell-types used in our analysis. Details of the tissues and cell-types found in the Tabula Muris dataset, and which ones we chose for analysis in this work are located in supplement table 6.

#### DRG scRNA-seq data

scRNA-seq data from the mouse DRG (Zeisel et al., 2018) was used to generate the transcriptome profile of individual cell-types within DRG. Expression values and metadata per cluster file (L5_All.agg.loom) provided by the original publication was used in our analysis. The NF1-3, NP1-6, PEP1-8, SATG1-2, SCHW cell-types from DRG tissue were selected for analysis. These were further grouped as described in the results section.

#### Enteric glial cell RiboTag RNA-seq data

Previously published RiboTag RNA-seq data of enteric glial cells from colon tissue was used (Delvalle et al., 2018). In this study, colonic inflammation was used to study how the translatome of enteric glia changes with inflammation. The RiboTag procedure driven by *Sox10*-cre was used to generate enteric glia translatome in vehicle treated and colonic inflammation conditions. Raw sequencing data were provided by the authors of the original paper, and mapped to gencode vM16 mouse genome annotation (Frankish et al., 2019) and quantified using STAR 2.6.1c (Dobin et al., 2013; Dobin and Gingeras, 2016). Reads per gene per sample were provided by STAR output and used in the downstream analysis.

#### Retrogradely traced neurons from the colon to DRG

Previously published scRNA-seq of DRG neurons retrogradely traced from colon (Hockley et al., 2019) were used in the interactome analysis for colonic inflammation. Reads per gene per cell, as well as the clustering and gene marker information were provided by the authors.

### Human Sequencing Resources

#### Human DRG sequencing

Human DRG tissue samples, previously sequenced and analyzed by our lab (Ray et al., 2018; North et al., 2019), were used for the transcriptome profile of hDRG. Normalized TPMs reported in those papers were used in the analysis here.

#### Synovial scRNA-seq from RA and OA patients

Kuo *et al. (Kuo et al., 2019)*, identified cell-types involved in different types of arthritis from scRNA-seq data of 940 human synovial CD14+ cells. One cluster was associated with Rheumatoid Arthritis (RA) samples when compared with Osteoarthritis (OA) samples (RA-enriched macrophages). Conversely, a second cluster was associated with OA patients when compared to RA patients (OA enriched macrophages). These 2 cell clusters were picked for the interactome analysis to compare macrophages from RA patients versus macrophages from OA patients. The clustering used in the original paper was provided by the authors and used in the analysis here. The original single-cell transcriptome data was provided as reads per gene per cell.

#### The Cancer Genome Atlas (TCGA) pancreatic cancer samples

Pancreatic cancer tissue sequencing data was acquired from TCGA database (Tomczak et al., 2015). Individuals that had both healthy tissue and cancerous tissue sequenced from the pancreas were selected for analysis in this paper. There were a total of 4 pairs of samples in TCGA pancreatic cancer database that fit the criteria and all pairs were used. Data were provided as normalized FPKMs and these numbers were used for downstream analysis.

### Reference lists for receptor and ligand pairs

Ramilowski *et al*. (Ramilowski et al., 2015) identified 2557 pairs of ligand-receptor interactions that were used as the basis for generating the full database of ligand-receptor pairs used here. In order to curate a more complete ligand-receptor list, we first collected gene lists from gene ontology (GO) databases, including HUGO (Yates et al., 2017) and the GO (Carbon et al., 2009) database. Genes under the GO terms of cell adhesion molecules (CAM), G-protein coupled receptor (GPCR), growth factor, ion channels, neuropeptides, nuclear receptors, and receptor kinases were collected. We then manually performed a literature search and used an existing database (Szklarczyk et al., 2019) to add additional ligand-receptor pairs. A total of 3098 pairs of ligand-receptor interactions were used in our interactome analysis (entire list shown in supplementary table 7). The COMPARTMENT database (Binder et al., 2014), or other literature sources for genes not found in that database, were used to define categories for ligands and receptors.

Not all ligand-receptor pair interactions are directly encoded by the genome so we added enzymes that are known to synthesize ligands to the database and paired these with receptors for the synthesized ligand. Some interactions between proteins found at the surface of cells do not have a clear ligand and receptor relationship, for instance some extracellular matrix and cell adhesion proteins. In these cases, we manually classified the receptor in the interaction as the gene that was expressed in the DRG.

### Statistics

#### Gene relative abundance normalization

The majority of the transcriptome datasets used in this paper were provided as raw read counts per gene per sample/cell, and were thus normalized to provide relative abundance measures for genes using standard approaches (Pachter, 2011). Reads per Million Mapped Reads (RPM) or udRPM (upper decile normalized RPM) were primarily used for further analysis. To keep the gene relative abundance consistent across different data sources, we calculated RPM for the enteric glial cells when we reanalyzed the dataset. Since we did not compare gene expression levels across different genes, normalization by gene length does not influence our analysis.

The human pancreatic cancer samples were provided as normalized Fragments per Kilobase per Million Mapped Fragments (FPKM) by TCGA, and RNA-seq data for human DRG samples were provided as normalized transcripts per million (TPM) by the original paper. These were also kept as their original relative abundance values for downstream analysis. Since these datasets were used for building different interactomes, different relative abundance measures for different datasets do not affect our analysis.

#### RPM

RPM of a gene was calculated with the following formula:

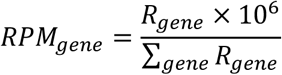

where *R*_*gene*_ stands for the read counts for a specific gene, ∑_*gene*_, *R*_*gene*_ stands for the read count sum of all genes.

#### udRPM

upper decile (ud) RPM was calculated to normalize sequencing depth and coverage differences across multiple samples. The formula used was:

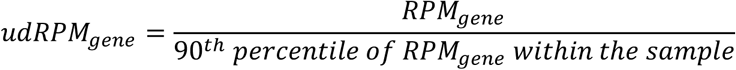

### Criteria for a gene to be considered as expressed in a given sample

#### Trinarization score

scRNA-seq data usually have low sequencing depth per cell. To better estimate if a gene is expressed, the trinarization score method was used. The trinarization score, as described in previous work by Zeisel *et al*. (Zeisel et al., 2018), was used to determine if a gene was considered detected in certain cell-types in specific tissues. The parameters used in the calculation of the trinarization score were: *f* = 0.05, α = 1.5, β = 2. Genes with probability P > 0.95 were considered expressed.

### Criteria for differential expression of gene relative abundance

#### Strictly standardized mean difference (SSMD)

Strictly Standardized Mean Difference (Zhang et al., 2006; Ray et al., 2019) was used to estimate effect size between 2 different conditions under comparison, using the following formula:

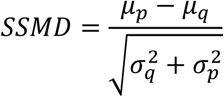

Where *p* and *q* stands for gene expression level across all cells in the first condition and second condition respectively, *σ* stands for standard deviation and *μ* stands for mean.

#### Bhattacharyya distance

The Bhattacharyya distance (Bhattacharyya, 1943; Wangzhou et al., 2020) was used to calculate the similarity of the probability distributions of scRNA-seq data from 2 different conditions using the following formula. The related Bhattacharyya Coefficient was used to calculate the amount of overlap in the area under the curve of the two sample distributions being compared:

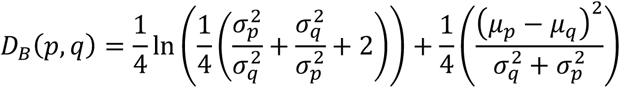

In this formula *D* stands for distance, *p* and *q* stands for gene expression level across all cells in the first condition and second condition respectively, *σ* stands for standard deviation and *μ* stands for mean. Genes with a distance > 0.3 between 2 conditions of scRNA-seq were considered differentially expressed.

### Interactomes

Each interactome was generated by intersecting the list of all genes and their expression levels in each tissue or cell type of interest with the ligand-receptor pair list described above (supplementary table 7). This resulted in a list of ligands, ligand expression levels, and the corresponding receptor gene names. The receptor genes were then intersected with the appropriate DRG RNA-seq dataset. This generated a resource which displayed the ligand gene name, ligand gene expression level in our tissue or cell-type of interest, receptor gene, and receptor gene expression level in the DRG. Each generated list was then filtered as described in detail below for each interactome.

### GO term analysis for ligands and receptors

To generate the GO term analysis for individual cell-types, the interactome analysis was performed between all 42 cell-types identified from Tabula Muris (Schaum et al., 2018) and the pooled PEP, NP and NF types of sensory neurons from the mouse DRG scRNA-seq dataset (Zeisel et al., 2018). Reads for each cell were pooled by the cell-types pre-identified in Tabula Muris dataset, and normalized to udRPM. For the interactome of individual cell-types, the udRPM were used to filter the ligand-receptor pairs. Any ligand-receptor pairs where the ligand expression level or the receptor expression level in all 3 DRG cell-types was less than 0.001 was considered ‘not detected’ and thus removed from the individual cell-type interactome. GO term analysis was then performed on the ligand and receptor genes for the individual interactomes using Enrichr (Chen et al., 2013; Kuleshov et al., 2016) and the top 5 (ranked by adjusted p-value) GO terms for GO biological process and GO molecular function were collected and shown in supplement table 2, sheets 0-42 as well as in Figure 1.

### Cell-type enriched interactome for 18 cell-type modules vs 3 types of sensory neurons

To identify gene-modules that were enriched in different cell-types, a cell-type and gene-module bi-clustering was performed on the udRPM per gene per cell-type matrix generated in the for the 42 cell type interactome. The scrattch.hicat package from the Allen Institute (Tasic et al., 2018) was used to generate clusters to identify gene modules. A factor of 1000 was multiplied by all numbers in the matrix to avoid a significant digit rounding impact on the result. Before clustering was performed, a log 2 transformation was performed on all 1000 x udRPMs with a smoothing value of 1 added to every number. The parameters for each round of clustering are listed in supplement table 8. Through iterative clustering, we identified a total of 8189 genes that were generically expressed across all pre-identified cell-types, and were used in the general interactome. Another 5117 genes were identified to have low to no expression across all cell-types and were excluded from the analysis. Twenty-five clusters of cell-types were identified in the fourth and final round of clustering, and were used in the cell-type enriched interactome.

scRNA-seq mouse DRG data (Zeisel et al., 2018) were pooled into 5 main groups: neurofilament (NF), non-peptidergic (NP), peptidergic (PEP), satellite glial cells (SATG), Schwann cells (SCHW). The mean of the normalized expression values was used for the expression level of each merged group, and udRPMs were calculated. The output from these calculations was then used as the transcriptome profile of different cell-types from the mouse DRG.

The interactomes for each cell-type cluster, as well as the general interactome, were then constructed as described above. Results were filtered by the receptor gene expression level in DRG sensory neurons. Ligand-receptor interactions with receptor expression level < 0.001 udRPM across all 3 types of sensory neurons (PEP, NF, NP) were excluded from the results.

### Interactome between different cell-types within the mouse DRG

Interactome analysis on mouse DRG cell clusters was performed in 2 separate directions: ligands from neuronal cells (NF, NP, PEP) signaling to receptors from glial cells (SCHW, SATG), and vice versa. All identified ligand-receptor interactions were filtered by whether the ligands and receptors were both expressed in at least 1 source cell-type. The trinarization score was calculated to determine if a gene was to be considered expressed in a certain cell-type (score > 0.95). Circle plots were generated to present these interactions using the Circos program (Krzywinski et al., 2009) where each identified ligand and receptor interaction is represented on the plot. To better present how certain groups of ligands or receptors interact with each other, the ligand and receptor genes on the circle plots were ordered by bi-clustering (Euclidean, average distance) of the ligand and receptor genes based on their interactions with each other. Additionally, each gene was identified as either NF, NP, PEP, SATG, or SCHW, and marked as such on the plot.

#### Interactome between retrogradely traced colonic sensory neurons and enteric glial cells in a colitis model

For the transcriptome of retrogradely traced colonic sensory neurons (Hockley et al., 2019), reads from all cells of the same cell-type were pooled together to generate the gene expression level per cell-type, and RPMs were calculated for further analysis. The interactome was then generated between these 7 cell-types and enteric glial cells after vehicle or DNBS treatment (Delvalle et al., 2018). Ligand-receptor interactions were filtered by the following criteria: (1) Ligand genes were required to be consistently expressed (>0.01 RPM) in enteric glial cells across all replicates in at least one condition (vehicle/DNBS treated); (2) Ligand genes were considered to be altered by treatment when the |*SSMD*| > *μ*_|*SSMD*|_ + 1.2*σ*_|*SSMD*|_; and (3) Receptor genes were required to be considered expressed using the trinarization score > 0.95. Interactions where the receptor genes were enriched in specific populations of colonic afferents are shown in Figure 5. The remaining ligand receptor interactions are provided in supplement table 4.

### Interactome between macrophages enriched in human RA synovial tissue versus OA, and hDRG

The interactome was generated between RA/OA enriched macrophages (Kuo et al., 2019), and human DRG (Ray et al., 2018). Ligand-receptor interactions were filtered by the following criteria: (1) Ligand genes were required to be considered expressed in either RA enriched macrophages or OA enriched macrophages using the trinarization score > 0.95; (2) Ligand genes were considered significantly differentially expressed between RA enriched macrophages and OA enriched macrophages, using Bhattacharyya distance > 0.3; and (3) Receptor genes were required to be consistently expressed (>0.1 TPM) in all 3 human DRG samples. Interactions where the ligand genes were highly expressed in RA enriched macrophages are presented in Figure 6. The interactions with ligand genes highly expressed in OA enriched macrophages are provided in supplement table 5.

### Interactome between human pancreatic cancer tissue and human DRG

The interactome was generated between 4 paired healthy/cancer tissue samples from pancreatic cancer patients (TCGA), and hDRG samples (Ray et al., 2018). Ligand-receptor interactions were filtered by the following criteria: (1) Ligand genes were considered significantly regulated in pancreatic cancer samples vs healthy samples by statistical testing (paired Student’s t-test, p-value < 0.05); (2) Receptor genes were required to be consistently expressed (>0.1 TPM) in all 3 hDRG samples. All interactions passing the filtering are presented in Figure 7 and 8.

## Acknowledgements

The authors would like to thank Drs. James Hockley and Ewan St. John Smith for help with the colonic single neuron sequencing data, Dr. Brian Gulbransen and lab for help with the enteric glia TRAP data and Dr. Zhenyu Xuan for clarifying TCGA metadata formats. We thank all the authors of the papers from which we used their sequencing data for their exemplary transparency in sharing the details of their work with us.

